# An RNA Virome associated to the Golden Orb-weaver Spider *Nephila clavipes*

**DOI:** 10.1101/140814

**Authors:** Humberto J. Debat

## Abstract

The golden orb-weaver spider *Nephila clavipes*, known for its sexual size dimorphism, is abundant and widespread in the New World. The first annotated genome of orb-weaver spiders, exploring *N. clavipes*, has recently been reported. The study, focused primarily on the diversity of silk specific genes, shed light into the complex evolutionary history of spiders. Furthermore, a robust transcriptome analysis provided a massive resource for *N. clavipes* RNA survey. Here, I present evidence of viral sequences corresponding to the first 10 extant virus species associated to *N. clavipes* and indeed, nephilids. The putatively new species are linked to ssRNA positive-strand viruses, such as *Picornavirales*, and to ssRNA negative-strand and dsRNA viruses. In addition, I detected sequence data of new strains of two recently reported arthropod viruses, which complemented and extended the corresponding sequence references. The identified viruses appear to be complete, potentially functional, and presenting the typical architecture and consistent viral domains. The intrinsic nature of the detected sequences and their absence in the recently generated genome assembly, suggest that they correspond to *bona fide* RNA virus sequences. The available RNA data allowed for the first time to address a tissue/organ specific analysis of virus loads/presence in spiders, suggesting a complex spatial and differential distribution of the tentative viruses, encompassing the spider brain and also silk and venom glands. Until recently, the virus landscape associated to spiders remained elusive. The discovered viruses described here provide only a fragmented glimpse of the potential magnitude of the *Aranea* virosphere. Future studies should focus not only on complementing and expanding these findings, but also on addressing the potential ecological role of these viruses, which might influence the biology of these outstanding arthropod species.

**Funding statement:** The author received no specific funding for this study.

**Ethics statements:** (Authors are required to state the ethical considerations of their study in the manuscript, including for cases where the study was exempt from ethical approval procedures)

## 1 Introduction

Recent advances in low-cost high-throughput metatranscriptomic sequencing is revolutionizing our understanding of the RNA virosphere. A considerable advancement by Li et al, (2015) depicted an unprecedented diversity of negative strand RNA viruses in arthropods. In addition, the same group (Shi et al, 2016) has recently reported more than 1.5 k RNA virus species infecting over 220 arthropods. These studies have shifted the paradigm of invertebrate virology, revealing a viral landscape phylogenetically and genomically diverse. Arthropods are an important source of virus diversity, and invertebrate virology is a flourishing and dynamic research field. In the past few years, the first spider infecting viruses have been described. Li et al (2015) identified seven species of ssRNA(-) viruses in pooled RNA samples encompassing five spider species: *Neoscona nautica*, *Parasteatoda tepidariorum*, *Plexippus setipes*, *Pirata sp*, and an unidentified *Araneae*. The viruses were assigned to the *Mononegavirales* order, two *Nairovirus* like and a *Plebovirus* like (*Bunyavirales*), an *Orthomyxoviridae* like and a new genus of non-segnented circular RNA viruses: *Chuvirus*. Remarkably, employing the same RNA library Shi et al (2016), pinpointed a ca. 80 collection of new virus species derived from spiders, corresponding to a wide range of RNA virus lineages, enriched in *Partitiviridae* and *Picornavirales* like viruses. In addition, Shean et al (2017) described six novel *Picornavirales* members from six different spider species found in Washington State. The viruses were identified in metagenomics libraries of the mygalomorph spider *Hexura picea*; the *Tetragnathidae* orbweaver *Metellina curtisi*; the triangle-weaving orbweaver *Hyptiotes gertschi;* the cobweb weaver *Theridion simile* and the crab spider *Xysticus cristatus*. The identified viruses presented low sequence similarity to reported picornaviruses (24%-47% by amino acid to the polyprotein) and phylogenetic analyses suggested they form a new clade within the *Picornavirales* order. The preceding studies have inaugurated *Araneae* virology by assessing the RNA landscape of only a dozen spiders. There are ca. 45.7 k spider species described, undoubtedly representing a massive reservoir of virus diversity, which remains unexplored.

The golden orb-weaver *Nephila clavipes* (Linnaeus, 1767) is a female-biased sexual-size dimorphic spider (Kuntner, 2009). It is a widespread and abundant species, distributed from southeastern United States to northern Argentina and from the Galapagos Islands to the Caribbean. They inhabit a broad range of habitats that vary from mild to strong seasonality (Higgins, 2000). *N. clavipes* spiders use golden-colored silks to spin orb webs. They are opportunistic predators, capturing diverse arthropods, and even small vertebrates (Higgins et al., 1992). Spider silks have a great potential for medical and industrial innovation, given their features of being both extremely strong and light (Agnarsson et al., 2010). *N. clavipes* generates a battery of silks derived from seven types of araneoid silk glands. This extensively studied species is considered the “ubiquitous workhorse of silk spider research” (Vollrath, 2000; Kaplan et al., 1993). Despite the importance of this orb-weaver spider, the molecular characterization of its genetic repertoire was lacking in the literature. Babb et al (2017) recently reported a sequencing *tour de force* that generated the first annotated genome of *N. clavipes*. Besides generating a genome assembly of 2.8 GB, the authors explored the RNA component of *N. clavipes* by multiple RNA-seq data from 16 different tissue/organ/individual isolates (whole body, brain, and silk and venom glands) collected from four female individuals. These 1.53 × 10^9^ reads corresponding to an assembly of 1.5 million transcripts, which represent the most extensive deposited RNA assembled data of any organism at the NCBI TSA database, were used as input for the objective of this study: The first identification and characterization of potential RNA viruses associated to *N. clavipes*.

## 2 Materials and Methods

In order to identify putative RNA virus associated with *N. clavipes*, the RNA data from Babb et al (2017) were integrated into *de novo* assembled transcriptomes generated for each isolate using strand-specific, ribosomal RNA (rRNA)-depleted, 100-bp paired-end reads. In sum, a total of 1,848,260,474 raw RNA reads were quality controlled and filtered, and the curated 1,531,402,748 reads were *de novo* assembled using Trinity (rel_2.25.13) yielding 1,507,505 unique strand-specific transcripts. These 1.53 × 10^9^ reads corresponding to an assembly of 1.5 million transcripts from Babb et al (2017) were used as input for virus discovery. In addition, the complete NR release of viral protein sequences was retrieved from https://www.ncbi.nlm.nih.gov/protein/?term=txid10239[Organism:exp]. The *N. clavipes* 1.5 million transcripts RNA assembly was assessed by multiple TBLASTN searches (max e-value = 1 × 10^−5^) using as probe the complete predicted non redundant viral proteins in a local server. Significant hits were explored by hand, and redundant contigs discarded. Potential virus genome segments sequences were curated by iterative mapping of reads using Bowtie 2 v2.3.2 http://bowtie-bio.sourceforge.net/bowtie2/index.shtml. To identify/rule out additional segments of no homology to the closely associated viruses I used diverse *in silico* approaches based on RNA levels by the sequencing depth of the transcript, predicted gene product structure, domain architecture, or conserved genome termini, and significant co-expression levels with the remaining viral segments. Potential open reading frames (ORF) were predicted by ORFfinder. Translated putative proteins were blasted against the non-redundant protein sequences NR database and best hits were retrieved. Predicted proteins were subjected to a domain-based Blast search against the Conserved Domain Database (CDD) v3.16 https://www.ncbi.nlm.nih.gov/Structure/cdd/cdd.shtml and integrated with SMART http://smart.embl-heidelberg.de/, Pfam http://pfam.xfam.org/ and PROSITE http://prosite.expasy.org/ to characterize the functional domains. Secondary protein structure was predicted with Garnier http://emboss.sourceforge.net/apps/release/6.6/emboss/apps/garnier.html, signal and membrane cues were assessed with SingnalP v4.1 http://www.cbs.dtu.dk/services/SignalP/ and prediction of transmembrane topology and signal peptides by Phobius http://www.ebi.ac.uk/Tools/pfa/phobius/. RNA secondary prediction was performed with the mfold Web Server http://unafold.rna.albany.edu/?q=mfold/rna-folding-form. Eventually, sequence similarity levels were visualized using the Circoletto tool with standard parameters http://tools.bat.infspire.org/circoletto/ Potential panhandle structures derived from partially complementary virus sequence termini were assessed with the RNAcofold web server http://rna.tbi.univie.ac.at/cgi-bin/RNAWebSuite/RNAcofold.cgi. Ribosomal frameshifting events were predicted using the KnotInFrame tool https://bibiserv.cebitec.uni-bielefeld.de/knotinframe. The potential functions of the ORFs products were predicted by the annotated data and similarity with known viral proteins. Virus RNA levels were calculated with Cufflinks http://cole-trapnell-lab.github.io/cufflinks/ or alternatively with the Geneious suite 8.1.9 (Biomatters inc.) as Fragments Per Kilobase of virus transcript per Million mapped reads (FPKM) based on million non-rRNA host transcriptome reads that map to the host genome assembly NepCla1.0. These normalized values avoid potential inconsistencies associated to the success of rRNA capture (RNA depleted libraries), the presence of other viruses and the diluting effects of variable non-host levels of reads. Tentative virus detections were contrasted on the *N. clavipes* NepCla1.0 genome assembly (NCBI accession no. GCA_002102615.1, 2.4Gb) by BLASTN searches (E-value = 1e10-6) and inspected by hand to rule out the identification of transcripts associated to putative integrated viruses or endogenous viral-like elements (EVE). In addition, the complete DNA derived raw read collection of Babb et al (2017) was mapped to the identified viruses to rule out unassembled EVEs. Amino acid sequences of the predicted viral polymerases or capsid proteins were used for phylogenetic analyses based on MAFTT 7.310 alignments http://mafft.cbrc.jp/alignment/software/ and unrooted FastTree maximum-likelihood phylogenetic trees http://www.microbesonline.org/fasttree/ with standard parameters. FastTree accounted for variable rates of evolution across sites by assigning each site to one of 20 categories, with the rates geometrically spaced from 0.05 to 20 and set each site to its most likely category by using a Bayesian approach with a gamma prior. Support for individual nodes was assessed using an approximate likelihood ratio test with the Shimodaira-Hasegawa like procedure. Tree topology, support values and substitutions per site were based on 1,000 tree resamples. Most sequence analyses results were integrated into the Geneious suite 8.1.9 (Biomatters inc.)

Based on sequence similarity to best hits, sequence alignments, predicted proteins and domains, and phylogenetic comparisons to reported species based in MAFFT and FastTree, I found evidence of 10 diverse new virus species and 3 new strains of reported virus species associated to *N. clavipes*.

## 3 Results and Discussion

### 3.1 *Nephila clavipes* Picornavirales like viruses

TBLASTN searches rendered sequences that could be linked to ssRNA positive strand viruses, most specifically to the *Picornavirales* order (Le Gall et al., 2008). These sequences were tentatively assigned to four putative new virus species. The proposed *Nephila clavipes* picorna-like virus 1 (NcPV1) is predicted to have a 10,198 nt long genome, presenting a single ORF between coordinates 761-10,147, encoding a putative 357.963 kDa and 3,128 aa long polyprotein. Domain prediction based on InterproScan, NCBI-CD database v3.15, THHM, PHOBIUS, SMART, Pfam, PROSITE and Garnier resulted in the identification of diverse motifs associated to *Picornavirales* coat and replicase proteins (RP) (Figure 1.A; Supp. Table 1). In addition, NcPV1 is similar to Wuhan spider virus 2 (WSV2) (Pairwise % Identity: 56.6% at the RNA level, and 39.5% at the predicted polyprotein level). WSV2 is currently unclassified, but tentatively assigned to the *Picornavirales*, within a newly proposed super clade of Picorna-Calici by the reporting authors (Shi et al., 2016). It is important to highlight that WSV2 was identified recently from a RNA pooled sample, a sequencing library made up by individuals of diverse classified and unclassified spiders: *Neoscona nautica* (14), *Parasteatoda tepidariorum* (3), *Plexippus setipes* (3), *Pirata sp*. (1), and 8 unrecognized individuals (*Araneae sp.*) (Shi et al., 2016). Thus, the specific host of WSV2 remains to be determined, even at the family level, which could contribute to the understanding of the evolutionary history of these viruses and their hosts.

**Figure 1.**
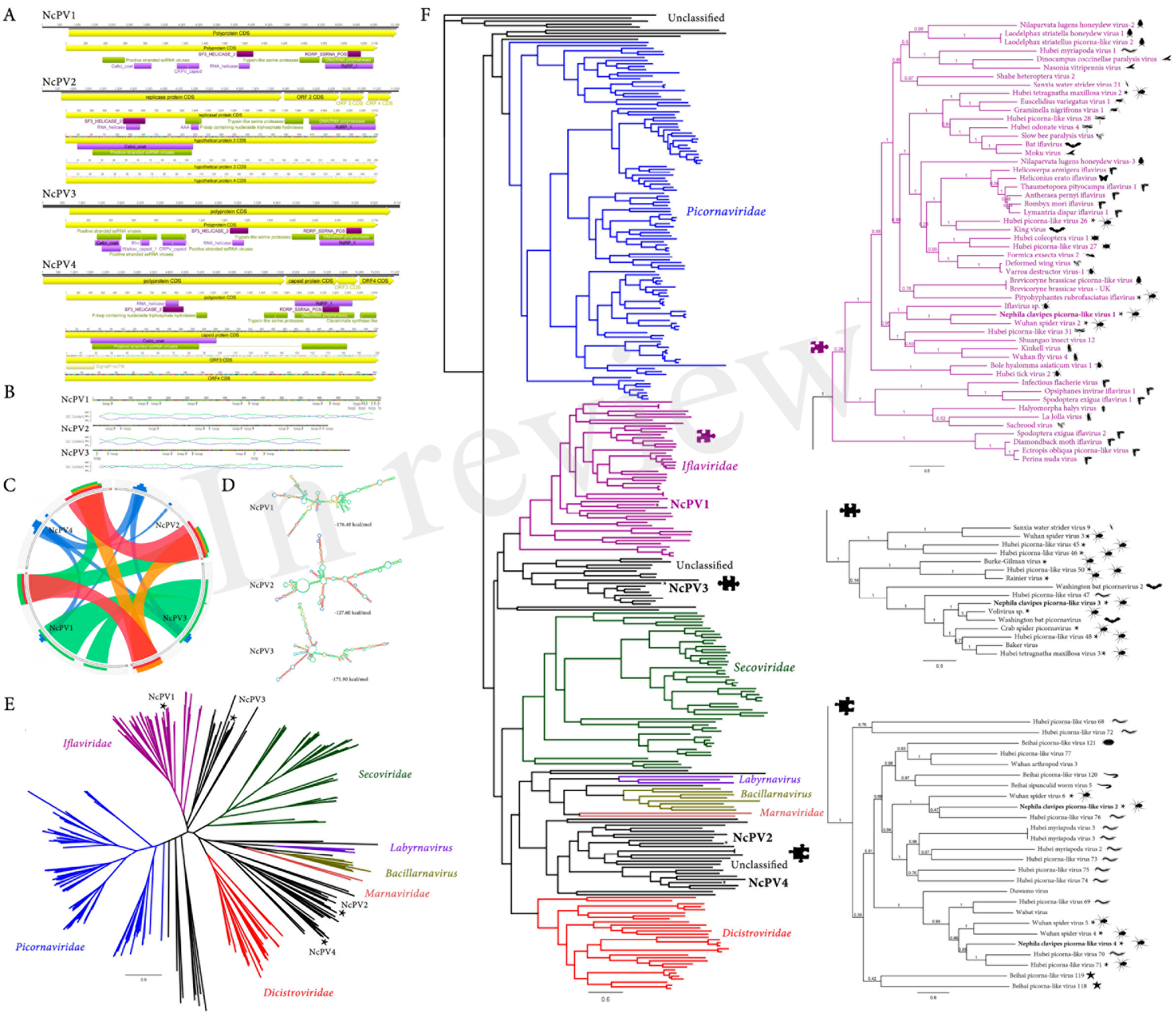
*Nephila clavipes Picornavirales* like viruses **A)** Genome graphs depicting genomes and predicted gene products of *N. clavipes* picorna-like viruses. Pfam, PROSITE and Superfamily predicted domains (E-value ≤ 1e-5) are shown in purple, bordeaux and green, respectively. Predicted domain data is available in Supp. Table 1-4. **B)** NcPV1-3 long A+U rich 5’ untranslated regions, presenting several UUUA motifs (loop) found typically in *Picornavirales*. p ercentage of GC and AU content are expressed in blue and green line graphs, respectively. **C)** Similarity levels of NcPV replicases expressed as Circoletto diagrams based on BLASTP searches with an E-value of 1e-1 threshold. RPs are depicted clockwise, and sequence similarity is visualized from blue to red ribbons representing low-to-high sequence identity. **D)** NcPV1-3 secondary structure of 5’ UTR, a RNA element predicted to function as internal ribosome entry site (IRES), allowing translation initiation in a cap-independent manner, as part of protein synthesis. **E)** Maximum likelihood unrooted branched phylogenetic tree based in MAFFT alignments of predicted replicase proteins of *Nephila clavipes* picorna-like viruses (black stars) and related viruses. Families of viruses of the *Picornavirales* order are indicated by colors. Scale bar represents substitutions per site. **F)** Rooted layout of the preceding phylogenetic tree. Magnifications of relevant regions of the tree are presented on the right and indicated by puzzle pieces. Reported hosts of viruses are represented by silhouettes. Branch labels represent FastTree support values. Complete tree showing tip species, host and virus assigned taxonomy labels are available as (Supp. Fig. 1-3). Abbreviations: NcPV1-4, *Nephila clavipes* picorna-like virus 1-4.

*Nephila clavipes* picorna-like virus 2 (NcPV2) shares the proposed Picorna-Calici ssRNA(+) super clade with NcV1, but presents a different genome organization and domain architecture (Figure 1.A, Supp. Table 2). NcPV2 presents an 11,699 nt RNA genome enclosing 4 putative ORFs. ORF1 (722-7,226 nt coordinates) encodes a putative RP of 274 kDa and 2,412 aa long. NcPV2 ORF2 (7,948-9,747 nt) encodes a putative 67.3 kDa coat protein, 599 aa long. ORF3 (9,792-10,562 nt) encodes a 27.7 kDa, 256 aa long protein similar to the hypothetical protein 3 of Hubei picorna-like virus 76 (E-value = 8e-14, 38% identity) and of Wuhan spider virus 6 (WSV6; E-value = 8e-14, 38% identity), both of unknown function. ORF 4 (10,702-11,457 nt) encodes a 29.7 kDa, 251 aa long protein, with a central coiled-coil region, and of unknown function. Although it is evident that NcPV1 and NcPV2 share several conserved domains, probably suggesting a common origin, their divergent spatial arrangement and their low sequence identity might be interpreted as an separation followed by recombination and reorganization of the putative proteins on the respective viruses. NcPV2 is similar to WSV6 (Pairwise Identity: 51.7% at the RNA level, and 28.5% at the predicted replicase protein). WSV6 was also recently identified from a pooled sample library of individuals of diverse spiders, and has been provisionally assigned to a Picorna-calici superclade. Its specific host spider remains unidentified.

An additional picorna-like virus could be associated to *N. clavipes*. NcPV3 is predicted to have a 9,141 nt long genome, presenting a single ORF between coordinates 680-8,914, encoding a putative 312.010 kDa and 2,744 aa long polyprotein presenting several *Picornavirales* associated domains (Figure 1.A; Supp. Table 3). The domain architecture of NcPV3 is equivalent to that of NcPV1, however their polyprotein similarity is low (20.6 % at the aa level), strongly suggesting that they are separate species. NcPV1-3 shared a characteristic long 5 ′UTR ranging from 620 to 760 nt. These particularly long UTRs could be associated to the cap-independent internal initiation of translation associated to picornaviral RNA, based on internal ribosome entry site (IRES) (Pilipenko et al., 1994). The 5′ non-translated region of NcPV1-3 resembles IRES by presenting a complex clover leaf like secondary structure, encompassing A+U rich hairpins, and presenting several UUUA loops found typically in *Picornavirales* (Figure 1.B and D; Pilipenko et al., 1989). Blastp searches using the RP indicate that NcPV3 shares 48 % similarity with the RP of Washington bat picornavirus (WBPV; E-value = 0.0; GenBank KX580885.1). WBPV was released to GenBank last year by the University of Washington Virology NGS group, and identified in a bat RNA library, but there is no related literature available. Interestingly, NcPV3 RP shares 41% similarity (E-value = 0.0) to Hubei tetragnatha maxillosa virus 3, which was reported by Shi et al (2016), and identified in a *Tetragnatha maxillosa* (*Tetragnathidae*) RNA library. This spider is a member of the *Araneoidea* superfamily, which includes the *Araneoidea* family, thus it is the closest spider to *N. clavipes* in which a virus was identified.

*Nephila clavipes* picorna-like virus 4 (NcPV4) shares the genome architecture of NcPV2. NcPV4 presents an 11,237 nt RNA genome enclosing 4 putative ORFs. NcPV4 ORF1 encodes a 2,533 aa long replicase protein (Figure 1.A; Supp. Table 4). NcPV4 ORF2 encodes a putative coat protein, 539 aa long similar to Wuhan spider virus 4 (WSV4) capsid protein (E-value = 1e-83, id 33%). NcPV4 ORF3 encodes a 202 aa protein similar to WSV4 hypothetical protein 3, of unknown function (E-value = 1e-20, id 49%). The 360 aa protein encoded in ORF4 shares no significant similarity to known proteins. NcPV1-NcPV3 and NcPv2-NcPV4 not only share genome organization, but also presents higher levels of similarity at their respective replicases, and sequence identity is significantly higher in equilocal regions suggesting a shared protein architecture (Figure 1.C). NcPV1-4 were explored by maximum likelihood phylogenetic trees derived from MAFFT alignments of their RPs, which showed that they cluster among several newly found picorna-like invertebrate viruses (Figure 1.E-F: Supp. Figure 1-3). Moreover, NcPV1-4 are separated in divergent sub-groups within this picorna-like clade. Even though the *N. clavipes* viruses are more closely related to unclassified *Picornavirales*, by generating phylogenetic trees based in assigned species I could hint some tentative affinities of the identified viruses with specific viral families. NcPV1 appears to be more closely related to the *Iflaviridae* family of *Picornavirales*, clustering among diverse invertebrate picornaviruses, and within a specific phylogroup of ticks, flies, odonata and spiders proposed picornaviruses, having as nearest neighbor the spider derived WSV2. NcPV3 clusters in a lineage of unclassified *Picornavirales* with affinity to the *Iflaviridae*. Interestingly, this new phylogroup, composed also by bat and centipede viruses, is highly enriched with spider derived viruses (11 of 16 viruses). NcPV2 and NcPV4 appear to be more closely related to a distinctive phylogroup of unclassified *Picornavirales*, which diverges branching between the *Dicistroviridae* and the *Marnaviridae* families of viruses. This phylogroup is enriched with *Myriapoda* derived viruses, however among NcPV2 and NcPV4 distinctive subgroups there are several spider derived viruses, such as WSV4, WSV5 and WSV6. There appears to be some distinguishing cues in the replicase sequences of arthropod picorna-like viruses, which is reflected in the particular clustering correlated with evolutionary history of putative hosts in phylogenetic trees. It is worth noting that these unclassified picorna-like viruses have been identified in broad non-targeted transcriptomic studies, and the biological and ecological implications of the virus presence remains unclear. The associated literature is exceptionally limited; thus until future studies explore the potential impact of these viruses on their host, it might be prudent not to speculate on their biology. Distant virus species of this order have been studied in more detail, such as *Acute bee paralysis virus* (*Picornavirales; Dicistroviridae; Aparavirus*) or *Sacbrood Virus* (*Picornavirales; Iflaviridae; Iflavirus*) which are reported to have drastic effects on their bee host, resulting in larvae death and sudden colony collapse (Govan et al., 2000; Ghosh et al., 1999).

### 3.2 *Nephila clavipes* Virgaviridae-like viruses

The putative *Nephila clavipes* virga-like virus 1 (NcVV1) is predicted to have a 10,569 nt long ssRNA (+) strand genome, presenting two partially overlapping ORFs, and three additional ORFs. ORF1 (9-6,809 nt coordinates) encodes a putative replicase protein (RP) of 260.4 kDa and 2,266 aa long (Figure 2.A; Supp. Table 5). ORF2 (6,736-8,088 nt coordinates), which is tentatively translated by ribosomal frameshifting, encodes a 51.8 kDa, 450 aa long hypothetical protein, sharing significant identity with the virion structural glycoprotein s2gp2 (E-value = 3e-05, 41% identity) corresponding to RNA 2 of the bisegmented *Chronic bee paralysis virus* (CBPV). To date, the ssRNA(+) CBPV remains unclassified by the International Committee on Taxonomy of Viruses (ICTV), and only its RP sequence presents similarities with members of the *Nodaviridae* and *Tombusviridae* families. It has been suggested that CBPV might be the prototype species of a new family of positive single-stranded RNA viruses (Olivier et al., 2008). ORF3 encodes a 177 aa protein presenting a SP24 domain of virion membrane proteins similar to the hypothetical protein 3 of Lodeiro virus (LV, E-value = 6e-33, id 41%). ORF4 and ORF5 encode a 423 and 102 aa hypothetical proteins of unknown function. NcVV1 shares with LV 51.8% nt sequence identity at the RNA level, and 28.4% at the RP protein. RP based phylogenetic trees, suggest that NcVV1 is further related to *Virgaviridae* distant like viruses, forming distinctive clusters of invertebrate viruses, such as the mosquito derived *Negevirus*. NcVV1 forms a divergent phylogroup with the proposed LV (Figure 2.B-C; Supp. Figure 4-6). LV is an unclassified virus that has been recently reported to be derived from the crab spider *Philodromus dispar* (*Philodromiae*) (Shean et al., 2017).

**Figure 2.**
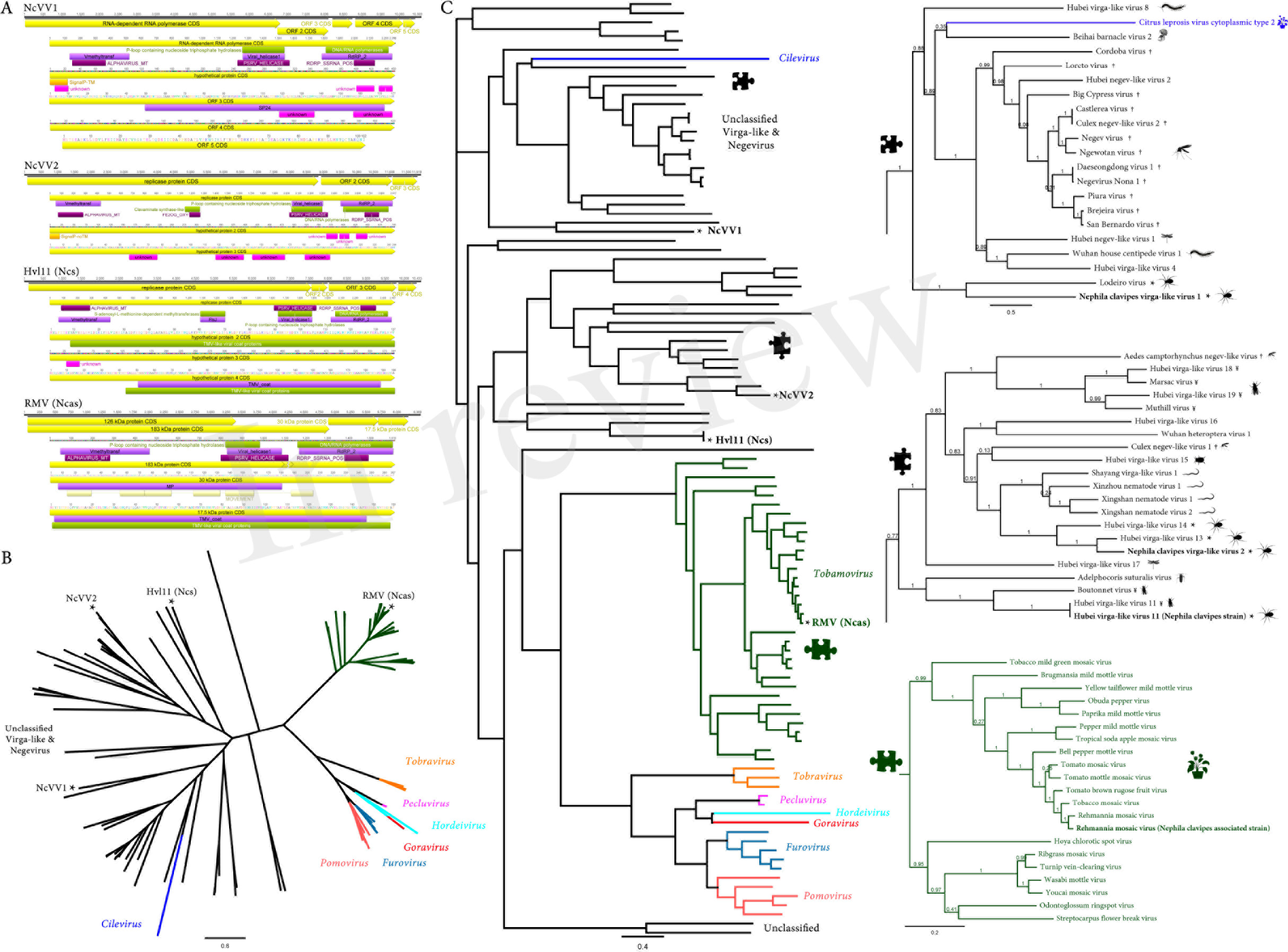
*Nephila clavipes Virgaviridae* like viruses **A)** Genome graphs depicting genomes and predicted gene products of *N. clavipes* virga-like viruses (NcVV1-2) and *N. clavipes* strains of Hubei virga-like virus 11 (Hvl11 (Ncs)) and *N. clavipes* associated strain of *Remania mosaic virus* RMV (Ncas)). Pfam, PROSITE, GENE3D, SignalP and Superfamily predicted domains (E-value ≤ 1e-5) are shown in purple, Bordeaux, pink, orange and green, respectively. Predicted domain data is available in Supp. Table 5-8. Abbreviations: WFV6-Ncs, *N. clavipes* strain of Wuhan fly virus 6; Hvl11-Ncs, *N. clavipes* strain of Hubei virga-like virus 11; RMV-Ncas, *N. clavipes* associated strain of *Remania mosaic virus*. **B)** Maximum likelihood unrooted branched phylogenetic tree based in MAFFT alignments of predicted replicase proteins of *Nephila clavipes* virga-like viruses (black stars) and related viruses. Genera of viruses of the *Virgaviridae* family are indicated by colors. Scale bar represents substitutions per site. **C)** Rooted layout of the preceding phylogenetic tree. Magnifications of relevant regions of the tree are presented on the right and indicated by puzzle pieces. Reported hosts of viruses are represented by silhouettes. † represent mosquitoes and ¥ flies. Branch labels represent FastTree support values. Complete tree showing tip species, host and virus assigned taxonomy labels are available as (Supp. Fig. 4-6).

*Nephila clavipes* virga-like virus 2 (NcVV2) shares with NcVV1 a super clade of Virga-like viruses. However the divergent NcVV2 clusters within a distinct group of ssRNA(+) viruses, some of them associated with nematodes, such as the recently reported Xinzhou nematode virus 1 (XzNV1) and Xingshan nematode virus 1 (XgNV1; Shi et al., 2016). Interestingly, the most closely related viruses based on replicase derived phylogenetic trees, correspond to two virga-like viruses derived from spiders: Hubei virga like virus 13 and 14 (Figure 2.B-C). NcVV2 presents an 11,919 nt ssRNA(+) genome enclosing three putative ORFs, of similar size and organization as XzNV1 and XgNV1. ORF1 (90-8,843 nt coordinates) encodes a putative replicase protein (RP) 334.9 kDa and 2,917 aa long (Figure 2.A; Supp. Table 6). NcVV2 ORF2 (8,934-11,051 nt) encodes a putative 80.7 kDa, 705 aa long hypothetical protein similar to the hypothetical protein 2 of Hubei virga-like virus 13 (E-value = 2e-108, 35% identity). ORF3 (11,084-11,818 nt) encodes a 27.3 kDa, 244 aa long protein, akin to the hypothetical protein 3 of XgNV1 (E-value = 1e-20, 32% identity).

### 3.3 *Nephila clavipes* bunya-like virus

The putative *Nephila clavipes* bunya-like virus (NcBV) is predicted to have a segmented genome. Sequence analyses resulted in the identification of two genome segments, tentatively assigned as genomes L and G. Genome segment L of NcBV is 7,365 nt long, presenting a single ORF between coordinates 56-7,285 nt (3′ to 5′orientation), encoding a putative 278.9 kDa and 2,409 aa long polyprotein (Figure 3.A; Supp. Table 9). Based on phylogenetic trees of the putative polyprotein, NcBV appears to be an ssRNA(-) virus, distantly related to the *Bunyavirales* order. NcBV predicted protein resembles the RdRP encoded typically in the Large genome segment of these multipartite viruses. Nevertheless, NcBV is more similar to several new “bunya-like RNA viruses” of diverse number of genome segments. Genome segment G of NcBV is 4,633 nt long, encoding a single 1,465 aa protein presenting a N-terminal integral transmembrane signal and a Nairovirus_M domain suggesting that this potential glycoprotein is associated to viral attachment and membrane fusion. Relaxed sequence TBLASTN searches based on Ns, Nm proteins of multipartite related *Bunyavirales*, co-expression retrieval of related transcripts, or domain based searches failed to retrieve any other potential gene segments of NcBV. L protein alignments of type species of the 9 *Bunyavirales* families and NcBV suggest a conservation of diverse motifs associated to the RDRP function of this protein (Figure 3.B). NcBV L protein presents an N-Terminal, Influenza-Like Endonuclease domain, essential for viral cap-dependent transcription (Reguera et al, 2010), and the functional motifs A-E of *Bunyavirales* polymerases (Kukkonen et al, 2005). *Bunyavirales* vRNAs present highly conserved, quasi-complementary 3′ and 5′ non-translated genome extremities, usually extending 13-19 nucleotides, which function as the promoter (Barr & Wertz, 2004). vRNA genome segments are packaged by multiple copies of the viral nucleoprotein together with the RdRP into filamentous ribonucleoprotein particles (RNPs), forming the functional replication and transcription units. The RNPs are circularized, by base pairing between the genome ends, forming a double-stranded “panhandle” mediating binding of both ends to the RdRP (Gerlach et al, 2015). NcBV L and G vRNAs present highly complementary genome termini, extending over 30 nt. In addition, RNA secondary predictions of potential duplex vRNAs suggest a stable, low FME structure supporting their putative role as promoter and tentative cues of *Bunyavirales* like replication (Figure 3.B). Phylogenetic trees of related virus sequences, suggest that NcBV might be a member of a new clade of invertebrate bunya- like divergent viruses, pivoting distantly to *Phasmaviridae*, *Nairoviridae*, and *Hantaviridae* (Figure 3.D-E; Supp. Figure 7-9). NcBV and related unclassified invertebrate bunya-like viruses cluster into a highly divergent clade which may eventually grant the proposal of a new family within the *Bunyavirales* order.

**Figure 3.**
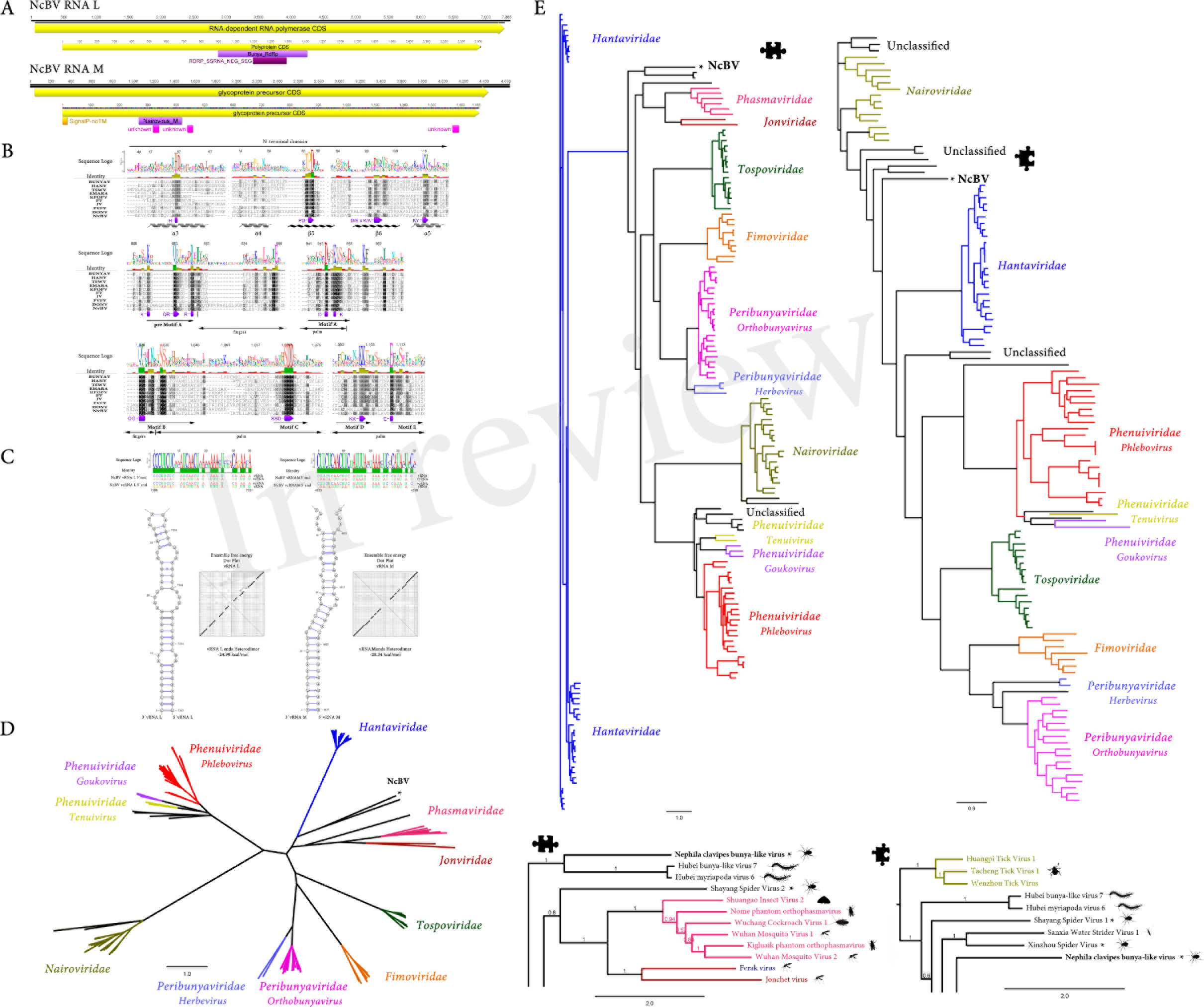
*Nephila clavipes Bunyavirales* like virus **A)** Genome graphs depicting genome segments and predicted gene products of *N. clavipes* bunya-like virus (NcBV). Pfam, PROSITE, GENE3D, and SignalP predicted domains (E-value ≤ 1e-5) are shown in purple, Bordeaux, pink and orange, respectively. Predicted domain data is available in Supp. Table 9. **B)** MAFFT alignment of L replicase protein of NcBV and type members of *Bunyavirales* families. Abbreviations: *Hantaan orthohantavirus* (*Hantaviridae* - HANV), *Europ*. *in mountain ah rin spoi-ussociated emaravirus* (*Fimoviridae* - EMARA), *Ferak orthoferavirus* (*Feraviridae* - FV), *Jonchet orthojonvirus* (*Jonviridae* - JV), *Dugbe orthonairovirus* (*Nairoviridae* - DONV), *Bunyamwera orthobunyavirus* (*Peribunyaviridae* - BUNYAV), *Kigluaik phantom orthophasmavirus* (*Phasmaviridae* - KPOPV), *Rift Valley fever phlebovirus* (*Phenuiviridae* - FVFV), *Tomato spotted wilt orthotospovirus* (*Tospoviridae* - TSWV). **C)** Sequence alignment of vRNA and vcRNA termini of NcBV genome segments L and G. Secondary structure prediction of RNA base pairing between the genome segment ends of NcBV, forming a double-stranded “panhandle” structure. NcBV L and G vRNAs present highly complementary genome termini, extending over 30 nt. Ensemble free energy of G and L vRNAs termini heterodimer expressed as dot plot. **D)** Maximum likelihood unrooted branched phylogenetic tree based in MAFFT alignments of predicted L replicase protein of *Nephila clavipes* bunya-like virus (black stars) and related viruses. Family of viruses of the *Bunyavirales* order are indicated by colors. Scale bar represents substitutions per site. **E)** Rooted layout of the preceding phylogenetic tree. Magnifications of relevant regions of the tree are presented below and indicated by puzzle pieces. Reported hosts of viruses are represented by silhouettes. Branch labels represent FastTree support values. Complete tree showing tip species, host and virus assigned taxonomy labels are available as (Supp. Fig. 7-9).

### 3.4 *Nephila clavipes Reoviridae*-like viruses

Sequence similarity searches resulted in the identification of numerous transcripts showing sequence identity to members of the *Reoviridae* virus family. *Reoviridae* are dsRNA, multipartite, non-enveloped, icosahedral capsid viruses (King et al, 2011). Based on sequence similarities with described viruses, co-expression levels and the pattern of presence or absence in diverse *N. clavipes* RNA libraries, and phylogenetic insights, the identified virus transcripts could be assigned to two tentatively new *Reovridae* like species. The proposed *Nephila clavipes* reo-like virus 1 (NcRV1) is predicted to have a multisegmented dsRNA genome. RNA segment 1 is 4,060 nt long, presenting a single ORF between coordinates 10-4,005 nt, encoding a putative 152.4 kDa, 1,331 aa long RP (Figure 4.A). NcRV1 RP appears to be highly divergent, BLASTP searches using the RP showed similarities to the RNA segment 1 of *Homalodisca vitripennis reovirus* (*Reoviridae; Sedoreovirinae; Phytoreovirus*) sharing a 23% aa identity (E-value = 3e-26). Hidden Markov Models (HMM) searches hint that the NcRV1 RP is similar to the Hubei reo-like virus 10 (HrlV10) RdRP (E-value = 4e-35), a recently described dsRNA *Reoviridae* like virus found in a *Odonata* (*Odonatoptera*) RNA pooled library (Shi et al., 2016). NcRV1 RNA segment 2 is 2,617 nt long, presenting a single ORF encoding a putative core protein of 93.7 kDa and 831 aa long. NcRV1 core protein is similar (E-value = 4e-11, pairwise identity 22%) to the putative minor capsid protein of Hubei reo-like virus 11. *Reoviridae* are multipartite viruses composed of ca. 10 dsRNA genome segments. Nevertheless, there are only two virus segments available for HrlV10 encoding a RP and a second RNA genome segment encoding a minor structural protein. Given the high divergence of NcRV1 with reported virus species, subsequent sequence analyses focused on co-expression levels of virus derived RNA levels, allowed to predict two additional tentative genome segments of NcRV1. Genome segment 3 of NcRV1 is 4,302 nt long, encoding a single 1,410 aa protein, similar to the hypothetical protein 4 of Hubei odonate virus 14 (Hov14, 23% similarity). Hov14 was identified in a dragonflies and damselflies pooled RNA library; thus its specific host remains unclear. Hov14 has been tentatively assigned to the reovirus-like superclade by Shi el al (2016). The NcRV1 genome segment 3 encoding protein is tentatively structural. Genome segment 4 is 1232 nt long encoding a single 365 aa protein, similar to the structural protein VP8 (26.1 % identity, E-value = 5.96e-02) of the *Reoviridae Banna virus* (BAV- *Seadornavirus*). BAV has been isolated from ticks and from humans suffering from meningoencephalitis (Attoui et al., 2000). VP8 is proposed to be the major virion protein of outer layer BAV viral particles, based on sequence similarity to VP6 of rotaviruses (Jaafar et al., 2005). An additional reo-like virus was detected in *N. clavipes. Nephila clavipes* reo-like virus 2 is composed of 10 dsRNA genome segments ranging between 1,096 and 3,837 nt long, generating a total genome length of ca. 23.6 kbps. Given the significantly low levels of NcRV2 vRNA in the *N. clavipes* libraries, genome segments were reconstructed by iterative mapping of reads to partial transcripts, resulting in a low coverage of the genome segments (ranging from only 12X to 20X). Segments 1 to 9 of NcRV2 segment presented a single ORF encoding a set of proteins which shares significant similarity (30% to 74% at the aa level) with the cognate proteins of Bloomfield virus, a proposed Reovirus, yet unclassified, described recently to be derived from a pooled sample of diverse wild caught *Drosophila* sp. flies (Webster et al, 2015). Additionally, segment 10 739 aa gene product shares similarity (E-value = 6.00e-37, id 29%) with the hypothetical protein encoded in RNA 3 of Hov14. Structural and homology based annotation rendered only a few signatures that could envision the functions of the divergent gene products of NcRV2. Homology detection and structure prediction with the HMM tool HHPred suggest that the gene products of RNA 2 and RNA 3 could have a structural function and that RNA 6 might be involved in methylation. Future studies could expand the current limited sequences reference set associated to these cluster of viruses, which might lead, in turn, to the eventual identification of highly divergent potential virus segments that should not be ruled out this soon, based on absence of evidence. RPs based phylogenetic trees hint that NcRV viruses form two divergent clusters within the *Reoviridae*, supporting their potential assignment to two new clades of invertebrate reo-like viruses (Figure 4.B-C; Supp. Figure 10-12). NcRV1 clusters into a divergent group of invertebrate viruses with affinity to the *Phytoreovirus* genus of plant infecting and insect vectored *Reoviridae*. Furthermore, NcRV2 is more closely related to another phylogroup of unclassified invertebrate viruses, linked to the *Fijivirus* genus of insect vectored, plant infecting *Reoviridae*. These incipient clusters of unclassified invertebrate viruses expand the diversity and evolutionary complexity within the *Reoviridae* family.

**Figure 4.**
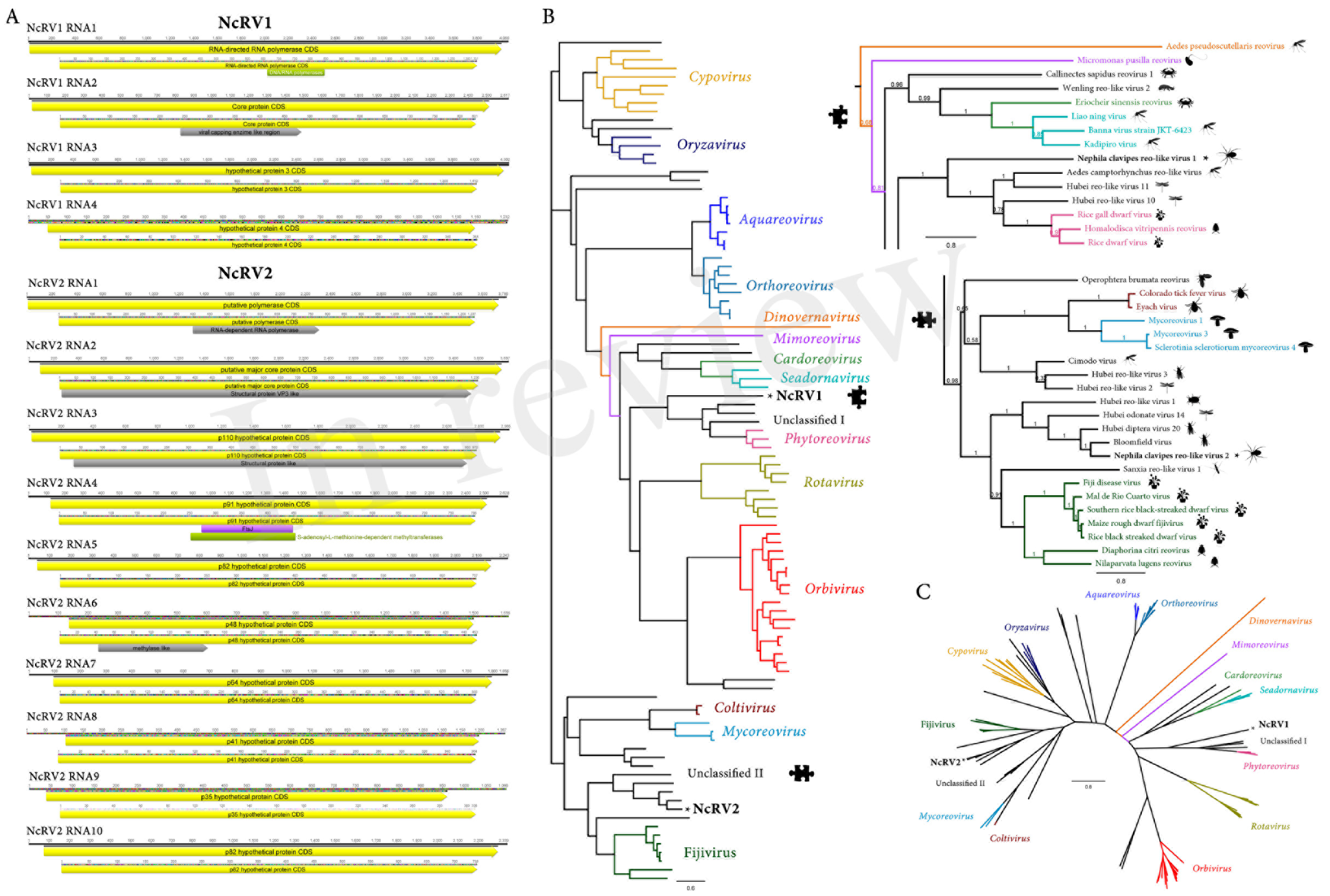
*Nephila clavipes Reoviridae* like viruses **A)** Genome graphs depicting genome and predicted gene products of *N. clavipes* reo-like virus 1 and 2 (NcRV1-2). Pfam and Superfamily predicted domains (E-value ≤ 1e-5) are shown in Bordeaux and green, respectively. Sequence regions presenting similarity with structural signatures are expressed in grey. **B)** Maximum likelihood rooted phylogenetic tree based in MAFFT alignments of predicted L replicase protein of *Nephila clavipes* reo-like viruses (black stars) and related viruses. Genera of viruses of the *Reoviridae* family are indicated by colors. Scale bar represents substitutions per site. **C)** Unrooted layout of the preceding phylogenetic tree. Magnifications of relevant regions of the tree are presented above and indicated by puzzle pieces. Reported hosts of viruses are represented by silhouettes. Branch labels represent FastTree support values. Complete tree showing tip species, host and virus assigned taxonomy labels are available as (Supp. Fig. 10-12).

### 3.5 *Nephila clavipes Astroviridae*-like virus

A highly divergent tentative virus transcript was identified in the *N. clavipes* RNA libraries. After further analysis involving genome architecture, gene product similarities with reported species, and phylogenetic insights based on a putative RdRP, I concluded that the sequence corresponded to a new species distantly related to the *Astroviridae* family, sharing some affinities also with the *Alphatetraviridae* family of viruses. *Astroviridae* were firstly described as electron microscopy detected particles with a “star-shaped” surface structure in stool samples from children with diarrhea. Astroviruses are spherical non-enveloped, 35-nm capsid with T=3 icosahedral symmetry, single-stranded RNA viruses (Monroe et al, 1993). Their monosegmented genome presents two partially overlapping ORFs that encode a protease and an RdRP (ORF1a-b) followed by a capsid precursor encoding ORF (ORF2) which is expressed as a subgenomic RNA (King et al, 2011). The *Nephila clavipes* astro-like virus (NcAV) presents a monopartite ssRNA(+) 7,569 nt long genome. NcAV presents two partially overlapping ORFs (ORF1a-b) followed by ORF2. Based on sequence analyses, ORF1a-b appear to encode a fusion protein of 1,756 aa generated by-1 ribosomal frameshifting (RF) (Figure 5.A). *Astroviridae* -1 RF is induced by a heptanucleotide “slippery sequence” of the form A_AAA_AAC, followed downstream by a RNA H-type pseudoknot structural element. The ribosome is stalled on the slippery sequence by the pseudoknot structure in 3’. In some cases, the ribosome would backtrack one nt, generating one mismatch between the cognate tRNA and the vmRNA. Thus, translation resolves on the backtracked ribosome in the-1 frame (Lewis & Matsui, 1995). Interestingly,-1 RF searches suggest that the corresponding slippery sequence of NcAV differs from that of *Astroviridae*, and is identical to the sequence of *Human immunodeficiency virus 1* (*Retroviridae*; *Lentivirus*) and *Sindbis virus* (*Togaviridae*; *Alphavirus*) (U_UUU_UUA). The heptanucleotide signal cue is supported by the presence of a highly stable H-type pseudoknot immediately downstream the slippery sequence (Figure 5.B). HHpred searches indicate that NcAV ORF1a presents a central serine peptidase region (with HDS catalytic triad) and a C-terminal zinc-binding motif/RNA binding suggesting that ORF1a is probably processed into at least three products (Figure 5.A; Supp. Table 10). ORF1b shares sequence similarity with Hubei leech virus 1 (HLV1 - E-value = 2.00e-16, id 29%), a proposed astro-like virus identified by Shi et al (2016). Intriguingly, ORF2 encodes a 648 aa capsid like protein, sharing sequence and structural similarity to the capsid protein of *Nudaurelia capensis omega virus* (NucOV), the type species of the *Omegatetravirus* genus of *Alphatetraviridae*. Interestingly, *Alphatetravirus* capsids present a T=4 icosahedral symmetry whereas *Astrovirus* capsids are T=3. NucOV coat protein assembles into a stable particle called the procapsid, which is 450 Å in diameter. Lowering the pH to 5.0 leads to a conformational change and maturation of the capsid, mediated by an autoproteolytic cleavage dependent on the presence of an Asn-570, which is at the cleavage site (Taylor et al, 2002). This residue is conserved in *Alphatetraviruses*, NcAV and two related unclassified viruses HLV1 and Hubei astro-like virus (Figure 5.D). The 3’ UTR of *Astroviridae* immediately adjacent to the poly(A) tail share a conserved secondary structure made up of three stem loops. This structure, reminiscent of the IRES *Picornavirales* region, is involved in virus replication (Monroe et al, 1993). NcAV presents a highly stable triple stem loop predicted RNA structure at its 3′ termini (Figure 5.C). Moreover, stem-I of NcAV presents at an equilocal region a UCUU motif. In *Human astrovirus 8*, stem-I UCUU mediates Polypyrimidine-Tract-Binding protein (PTB) binding to the 3’ UTR, which is required for Astrovirus replication (Espinosa-Hernández et al, 2010). RP and CP phylogenetic trees were generated to gain insights about the potential taxonomy of this divergent virus (Figure 5.E-F; Supp. Figure 13-15). RP based trees suggest that NcAV is distantly related to *Astroviridae*, clustering in a distinctive clade of invertebrate derived viruses, extensively separated from the mammal (*Mamastrovirus*) and bird (*Avastrovirus*) infecting *Astroviridae*. CP based trees add some complexity to the picture (Figure 5.F). NcAV also clusters within a new group of invertebrate viruses, but it appears to branch more closely to *Alphatetraviridae, Permutotetraviridae* and *Sinaivirus* than to *Astroviridae*. Future studies should complement the results reported here, which may lead to a better understanding of the evolutionary history and taxonomical assignment of NcAV and related viruses.

**Figure 5.**
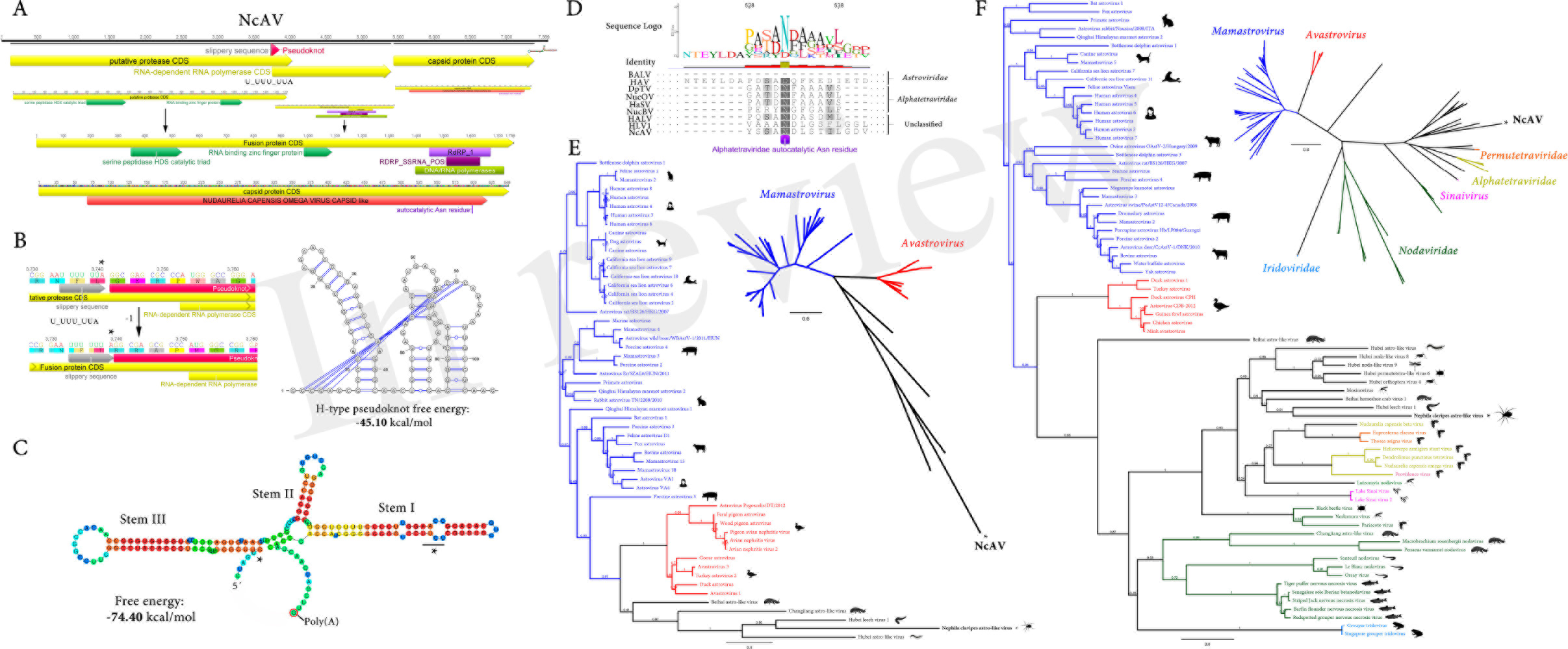
*Nephila clavipes Astroviridae* like virus **A)** Genome graphs depicting genome and predicted gene products of *N. clavipes* astro-like virus (NcAV). Pfam, PROSITE and Superfamily predicted domains (E-value ≤ 1e-5) are shown in purple, bordeaux and green, respectively. Functional domains determined by HHPRED are indicated in dark green. Predicted domain data is available in Supp. Table 10. **B)** NcAV predicted-1 ribosomal frameshifting is induced by a heptanucleotide “slippery sequence” of the form U_UUU_UUA, identical to the form of HIV1 (*Retroviridae; Lentivirus*) and *Sindbis virus* (*Togaviridae; Alphavirus*), followed downstream by a highly stable RNA H-type pseudoknot structural element. **C)** NcAV presents a highly stable triple stem loop predicted RNA structure at its 3’ termini. Stem I of NcAV presents at an equilocal region a UCUU motif (black star) which in *Human astrovirus 8* mediates PTB binding to the 3’ UTR, required for Astrovirus replication. **D)** MAFFT alignment of the capsid like protein of NcAV and related *Astroviridae, Alphatetreviridae* and unclassified astro-like viruses. Maturation of the capsid in alphatetraviruses is mediated by an autoproteolytic cleavage dependent of the presence of an Asn-570, which is at the cleavage site, and is present in NcAV and *Alphatetraviridae* and unclassified astro-like viruses, but not in *Astroviridae* viruses. Abbreviations: Beihai astro-like virus (BALV), *Human astrovirus* (HAV), *Dendrolimus punctatus tetravirus* (DpTV), Nudaurelia capensis omega virus (NucOV), Helicoverpa armigera stunt virus (HaSV), Hubei astro-like virus (HALV), Hubei leech virus 1 (HLV1). **E)** Maximum likelihood unrooted branched phylogenetic tree based in MAFFT alignments of predicted RP protein, or Capsid protein **(F)** of *Nephila clavipes* astro-like virus (black stars) and related viruses. Genera of viruses of the *Astroviridae* family and related virus families are indicated by colors. Scale bar represents substitutions per site. Rooted layout of the preceding phylogenetic trees are also presented. Reported hosts of viruses are represented by silhouettes. Branch labels represent FastTree support values. Complete tree showing tip species, host and virus assigned taxonomy labels are available as (Supp. Fig. 13-15).

In addition to the tentative new virus species detected, new strain of reported invertebrate virus associated to *N. clavipes* were identified and are presented as Supplementary Data 1. It is worth mentioning that every detected potential virus sequence was assessed on the whole genome assembly of *N. clavipes* (NepCla1.0; GenBank accession GCA_002102615.1; Babb et al., 2017) and no evidence that the tentative viruses could be derived from integration of virus-related sequences into the genome were found. Moreover, the complete collection of raw DNA sequencing data, ascending to a total of 4,094,217,472 read pairs were mapped to the identified viruses to explore the potential presence of unassembled virus like DNA. No virus like sequences were detected on the raw data of Babb et al., (2017). Furthermore, the facts that the detected virus sequences corresponded to the full length of the putative virus, present unaltered encoding and spacer regions, and maintain the typical domain architecture of related viruses, support the assumption that the identified sequences correspond to *bona fide* extant *N. clavipes* viruses. Moreover and importantly, in the case of multi-segmented nature viruses (*Partitiviridae*, *Reoviridae*), the corresponding and expected segment RNAs were found as independent units, presenting expected UTRs, ORFs and predicted gene products and shared a dynamic pattern of presence/absence and co-expression levels in the diverse RNA libraries.

### 3.6 Tissue/Organ presence and RNA levels of *N. clavipes* viruses

The *N. clavipes* multiple RNA-seq data derived from 16 different tissue/organ/individual isolates (whole body, brain, and individual silk and venom glands) collected from four female individuals (Babb et al., 2017) allowed for the first time to address the presence of a spider virus RNA at the tissue/organ level (Figure 6; Supp. Figure 21; Supp. Table 11). Virus levels were expressed as FPKM, mean coverage calculated, variants/polymorphism estimated among samples, and the tentative virus sequences were curated based on base frequency. Virus transcript presence and levels were complex, consistent, and varied by species, individuals and tissue/organs assayed. A total 5,286,563 absolute reads were assigned to be derived from viruses. RNA differential accumulation could be associated to specific virus species independently of taxonomy associations. For instance, in terms of absolute virus reads NcPV2 and NcRV1 accumulated at high levels (a total of 188.7 and 151,2 million nt among samples), while NcPV4 and NcRV2 accumulated at low viral titers (1.35 and 0.33 M nt; Figure 6.A). Essentially, virus derived RNA were retrieved on every sample, and to my knowledge this is the first time that a virus derived nucleic acid is detected specifically in silk and/or venom glands, and in the brain of spiders (Figure 6.B). If virus presence is estimated at the individual spider level, NcPV1 and NcPV2 were conclusively detected in every female spider sample, at different transcript levels, which varied in relation with the sampled organ (Figure 6.C-D, F). NcAV was detected in most spiders but not in Nep-7. NcVV2 and NcBV were detected in both Nep-7 and Nep-9 (Figure 6.D,F). HvlV11 (Ncs) was detected in Nep-8 and Nep-9, WFV6 (Ncs) only in Nep-9 and RMV (Ncas) only in a specific silk library of Nep-9 (Figure 6.C). Notably, the detection and RNA levels of multipartite predicted viruses (NcRV1, NcRV2, NcBV and WFV6) were consistent for the corresponding RNA genome segments among samples, independently confirming virus presence on selected libraries (Figure 6.D, G). It is important to highlight that the RNA virus estimated loads differed significantly among samples. For instance, NcPV1 levels were relatively high among most tissue samples but not on brain samples. On the contrary, NcRV1 levels were relatively low among samples, but strikingly spiked specifically on brain tissues, as is the case for NcPV2 on the Nep-8 sample. In general, as a whole, the presence of virus RNA was significantly accumulated in additional magnitude at the brain tissue, ascending in one sample to a striking ca. 6.1 % of total detected RNA reads (Figure 6.D, E). The biological significance of this finding, though interesting, remains unclear. It is tempting to associate over accumulation of virus RNA levels in the brain with central nervous system (CNS) immune privilege, a phenomenon widely studied in mammals (Carson et al,. 2006). A potential viral neurotropism could be linked to eventual behavioral altered phenotypes on infected individuals, as is the case for rabies in mammal hosts (Tsiang et al., 1983). Virus tropism literature on arthropods is scarce, but there are a few studies exploring tissue specific viruses and its implications. For instance, a picorna-like virus associated to a braconid wasp parasite, the *Dinocampus coccinellae* paralysis virus (DcPV), replicates in the host’s nervous tissue and induces a severe neuropathy. Interestingly the wasp transmits the virus to a coccinellid host, the Spotted lady beetle *Coleomegilla maculate*, in which virus replication induces an antiviral immune response that correlates with paralytic symptoms. This behavior manipulation of the coccinellid, characterized by tremors, gait disturbance and limited movements, facilitates parasitism and correlates with virus RNA levels in the cerebral ganglia (Dheilly et al., 2015). This remarkable phenomenon, which Stilling et al (2016) suggest as an example of virus driven puppeteers of neural function and behaviour, a kind of brain’s Geppetto, appear not to be incidental. The putative neurotropism correlated to neurological symptoms of DcPV has been reported for other CNS accumulating Picornaviruses, which are implicated in severe paralytic symptoms on their arthropod hosts. This is the case of the CBPV and of the *Cripavirus Aphid Lethal Paralysis Virus* (Williamson et al., 1988). It has been suggested that the potential behavioral alteration of virus host could be induced by the parasites to enhance virus replication and transmission, or a response of the host to avoid spread of infection. Host manipulation by behavioral alteration could have wide evolutionary and ecological importance, given the high prevalence of viruses among invertebrates (Han et al., 2015). Interestingly, the higher replication of DcPV in heads has been correlated with a transient downregulation of several genes involved in the antiviral response such as Toll 7 and PI3K (antiviral autophagy) and importantly of Dicer2, Ago2, R2D2 and C3PO (antiviral RNA interference). Therefore, a reduced RNAi activity at arthropods CNS, which is the primary and most important antiviral immune response in insects such as Drosophila and mosquitoes (Galiana-Arnoux et al., 2006; Blair 2011), could be linked to an eventual spike in virus RNA levels in CNS. Salazar et al., (2007) report a dynamic virus presence in midgut and salivary glands. They mention that the mosquito nervous system (used as control) presented large and persistent amounts of Dengue virus type 2 antigens. Future studies should assess whether my spider results are specific an anecdotal or if arthropod viruses are in fact over accumulated in nervous system tissues. Furthermore, the diverse viruses were consistently detected in the independent silk samples corresponding to the same spider, at a similar FPKM level, suggesting the accumulation of viruses is more-less steady among the diverse silk glands. Interestingly, virus RNA was also found in venom glands, NCPV1 being the most abundant. Female *Diachasmimorpha longicaudata* parasitic wasps are associated with two vertically transmitted RNA viruses that are present in the host venom glands, D. longicaudata entomopoxvirus and D. longicaudata rhabdovirus (Simmonds et al., 2016). Given the antecedent of the parasitic wasp *Dinocampus coccinellae* (Dheilly et al., 2015) it is tempting to suggest that the accumulation of virus in venom glands in *D. longicaudata* could be associated with the parasitic process. In *N. clavipes* the evolutionary implications of virus accumulation in venom glands is unclear. It would be interesting to explore if there could be some association with virus presence and predatory behavior or outcome. In addition, when whole spiders were sampled, the detected viruses were found to be at lower RNA levels than in the specific silk/venom/brain libraries (Figure 2.E-F). Although in the context of a small sample size and the fact that the whole body libraries derive from different sampled individuals than the tissue libraries, I cautiously speculate that perhaps these specific sampled tissues are enriched on RNA virus loads. More complex distribution/presence absence patterns are available in Supp. Figure 21 and Supp. Table 11.

**Figure 6.**
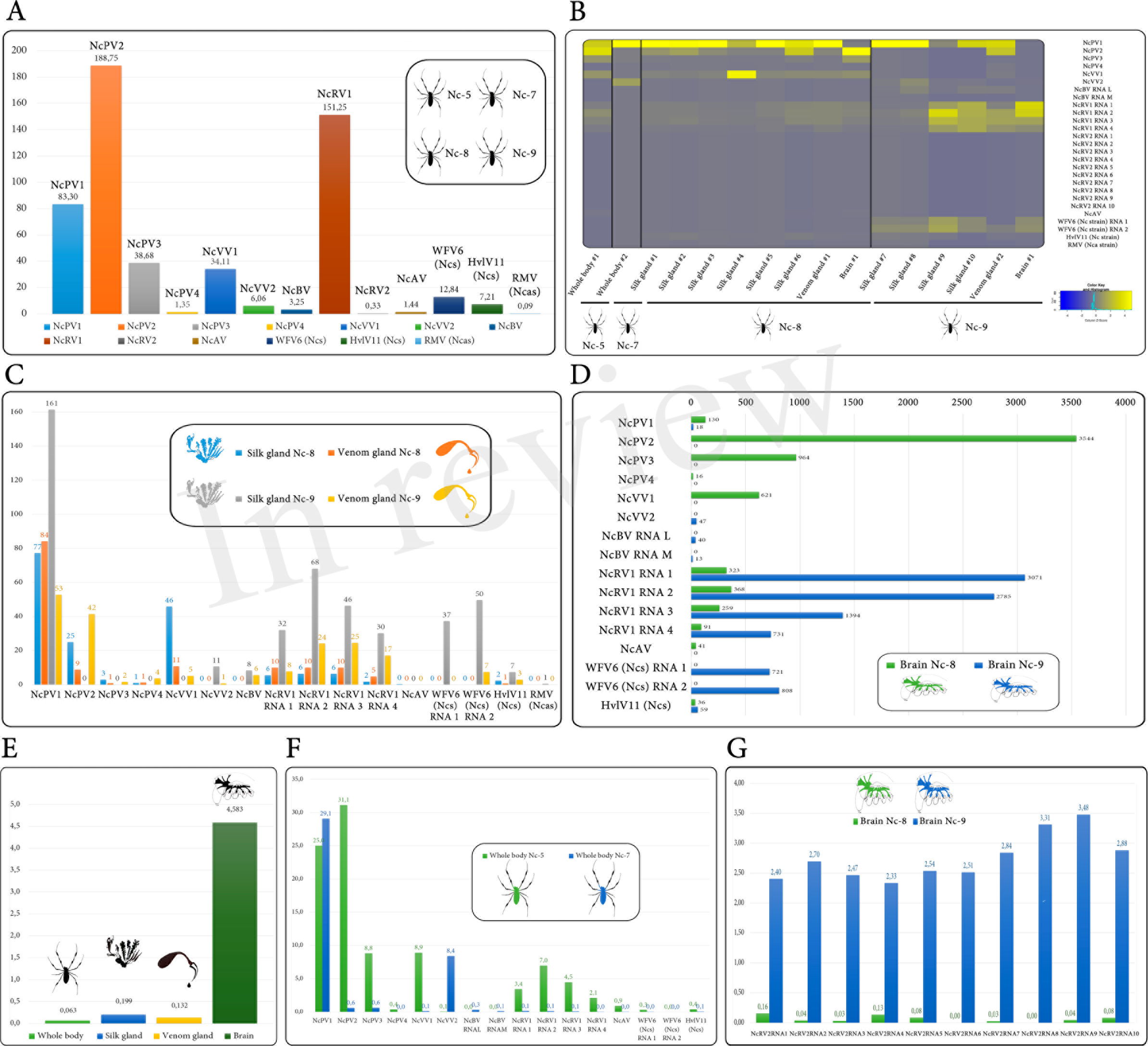
Graphs bars and heatmap describing virus RNA transcript levels assayed in two whole body spider samples, ten individual silk glands, two venom glands, and two brain isolates collected from four females, Nep-5, Nep-7, Nep-8 and Nep-009. Values are expressed as FPKM, where M indicates million of non-rRNA host transcriptome reads that map to the host genome assembly NepCla1.0 in **B, C, D, F** and **G**, and as total virus derived million nt in **A**, or non-rRNA host transcriptome percentage reads that map to viruses in **E**. FPKM Values corresponding to each sample are available as Supp. Table 11.

**Table 1.**
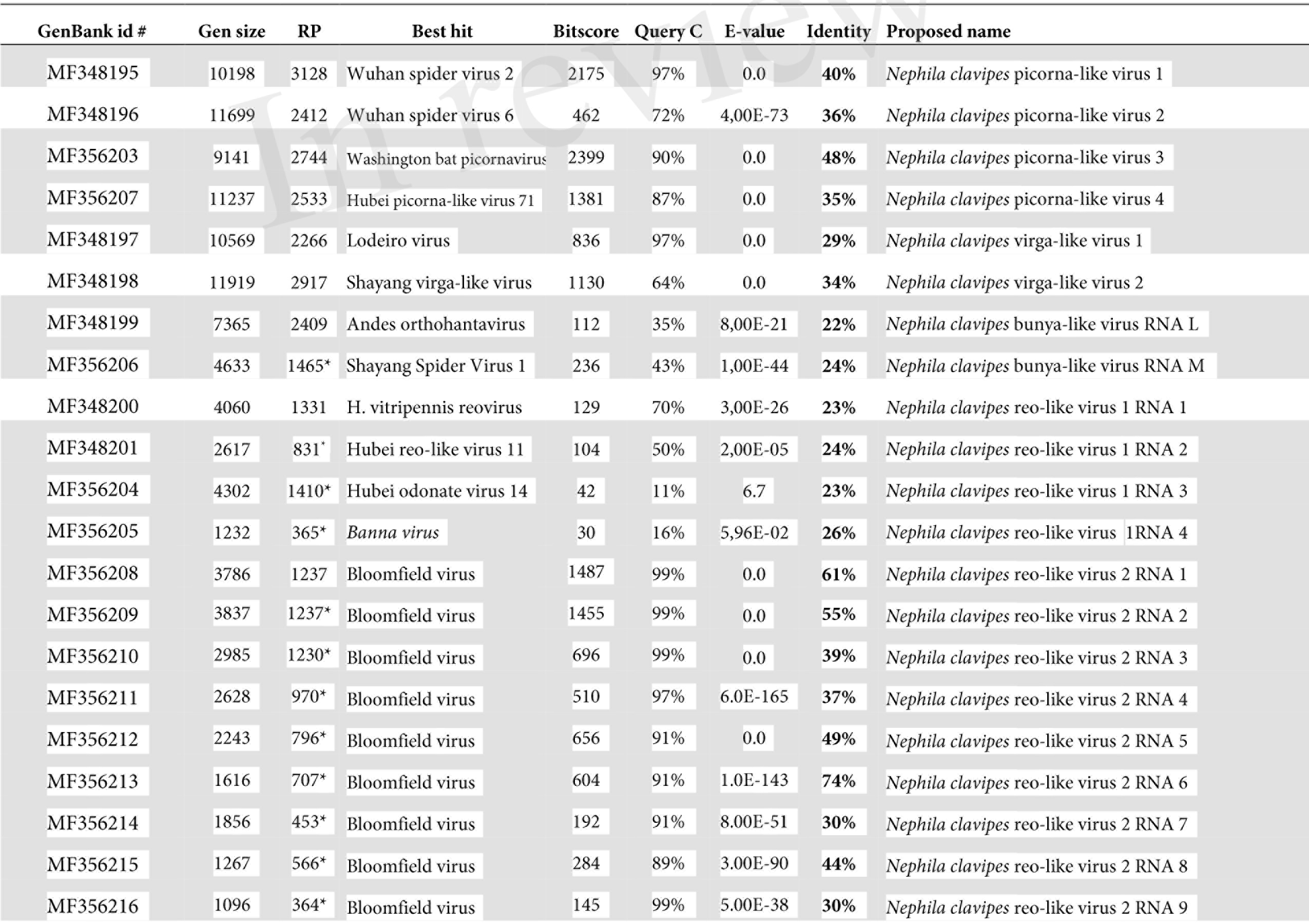
*Nephila clavipes* viruses, genome composition, similarity to reported viruses and proposed names.

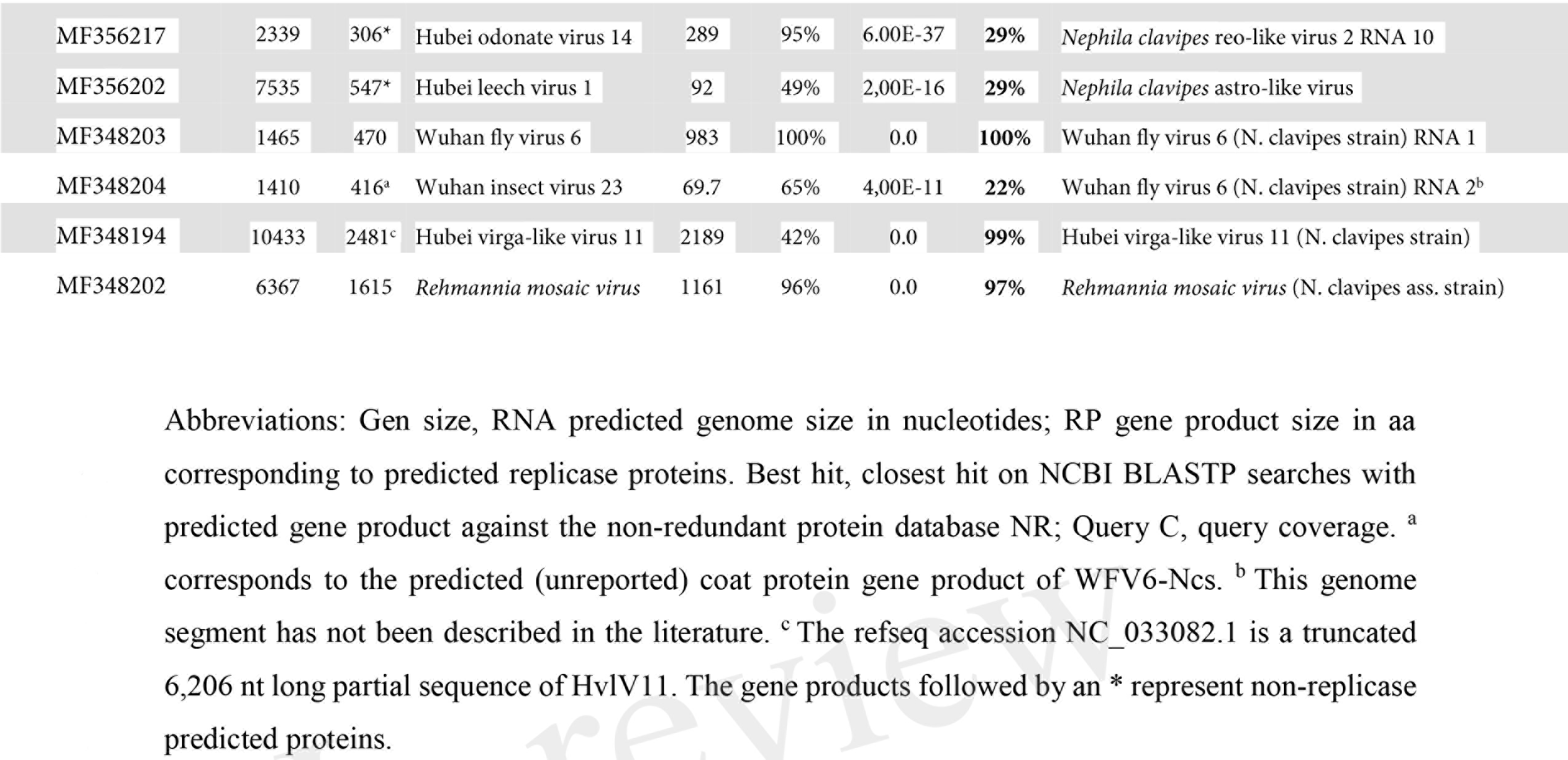

## 4 Conclusions

Regardless of sample size, and the limited number of detected viruses, it is interesting to highlight that most detected sequences corresponded to new unreported virus species. Spiders could be an important reservoir of viral genetic diversity that ought to be assessed. Widespread consistent RNA accumulation of diverse putative viruses on independent profiled samples, sequence structure and domain architecture, supports the assumption that the identified sequences correspond to *bona fide* viruses. It is not easy to speculate about the biological significance of the presence, accumulation, and distribution of these potential viruses in the context of limited literature. The brain enrichment of RNA virus loads appears not to be incidental, and could be associated to a potential effect on the spider host. The accumulation of viral RNA on silk and venom glands may have some evolutionary relation with virus horizontal transfer. Future studies should focus not only on complementing and expanding these findings, but also on addressing the potential ecological role of these viruses, which might influence the biology of these outstanding arthropod species.

## 5 Data availability

*Nephila clavipes* associated virus sequences have been deposited in NCBI GenBank (Accession numbers MF348194 to MF348204). Data from Babb et al (2017) are available through the central BioProject database at NCBI under project accession PRJNA356433 and BioSamples accessions SAMN06132062-SAMN06132080. All short-read sequencing data are deposited in the NCBI Short Read Archive (SRX2458083-SRX2458130), and transcriptome data are available at the Transcriptome Shotgun Assembly (TSA) under accession GFKT00000000.

## 7 Author Contributions

HJD designed the study, conducted all bioinformatics analysis, interpreted the data, wrote and approved the final manuscript.

## 8 Acknowledgments

I would like to express sincere thanks to Dr. Benjamin Voight for his encouragement in communicating these findings, which are only possible thanks to the remarkable work of his team and colleagues. Additional thanks to Dr. Casey Greene and colleagues for taking from words to action the active support and promotion of secondary analysis of data, which could redound in new discoveries and hypotheses, and a paradigm shift in research practices. Special thanks to Dr. Max L. Nibert for helpful discussions and insightful comments.

